# CDK4/6 inhibition sensitizes breast cancer to NK cell therapy by inducing immune-interactive surface proteins

**DOI:** 10.64898/2026.04.19.719504

**Authors:** Yinchong Wang, Evdokiya Reshetnikova, Nar B. Katuwal, Vijaya Bharti, Marcelo S F Pereira, Bridget A Oppong, Dean Lee, Arjun Mittra, Aharon G. Freud, Anna E. Vilgelm

## Abstract

CDK4/6 inhibitors are standard-of-care for metastatic estrogen receptor-positive (ER+) breast cancer, yet the development of resistance remains a significant clinical hurdle. While CDK4/6 inhibitors are primarily recognized for their ability to induce cytostasis, their role in modulating innate immune responses remains poorly defined. Here, we demonstrated that CDK4/6i treatment remodels the tumor cell surface to favor recognition and elimination by Natural Killer (NK) cells. Using a diverse biobank of patient-derived organoids (PDOs), we found that CDK4/6 inhibition robustly upregulated the adhesion molecule ICAM-1 and the NKG2D stress ligands (ULBP2/5/6 and MICA/B). This NK-engaging cell surface phenotype was driven by a bifurcated signaling network: NF-κB signaling orchestrated ICAM-1 induction, while the PI3K/mTOR pathway regulated the expression of stress ligands. Functional assays confirmed that these ligands were indispensable for NK cell-mediated elimination of breast cancer cells. *In vivo* studies using ER+ PDX models revealed that a brief seven-day primer treatment with the CDK4/6 inhibitor abemaciclib was sufficient to sensitize tumors to NK cell therapy, significantly inhibiting tumor growth and prolonging survival. We also observed efficacy with a concurrent dosing strategy that delayed the onset of acquired resistance. These findings provide a mechanistic rationale for combining CDK4/6 inhibitors with NK cell therapy. This “prime and kill” approach offers a promising strategy to overcome therapeutic resistance and improve outcomes for patients with metastatic ER+ breast cancer.

## INTRODUCTION

Breast cancer (BC) is the most common malignancy and the second leading cause of cancer-related mortality among women in the United States (*1*). Hormone receptor (estrogen receptor (ER) and/or progesterone receptor (PR))-positive and HER2-negative breast cancer (HR+ HER2-BC) is the most prevalent subtype. Approximately 25% of HR+ HER2-BC patients eventually progress to metastatic disease, which remains the primary cause of death in this population (*2, 3*). Currently, the frontline standard of care for metastatic disease involves endocrine therapy (ET) combined with cyclin-dependent kinase 4/6 inhibitors (CDK4/6i), such as abemaciclib, ribociclib, or palbociclib. While these combinations show profound initial efficacy by inducing cell-cycle arrest, they do not typically eliminate tumors, and patients inevitably develop resistance (*4*). This clinical reality underscores an urgent need for therapeutic strategies that move beyond cytostasis toward complete tumor elimination.

The canonical mechanism of CDK4/6i involves blocking the phosphorylation of the retinoblastoma 1 (Rb1) protein, thereby preventing E2F-mediated transcription and halting cells in the G1 phase of the cell cycle (*5, 6*). However, plentiful evidence suggests that these drugs possess significant immunomodulatory properties. For instance, prolonged cell cycle arrest induced by CDK4/6i treatment can lead to a damaged cell state, known as cellular senescence, characterized by stable cell cycle arrest and the secretion of numerous immune-related proteins, including pro-inflammatory cytokines (*7–12*). We reported that CDK4/6i increased the production of inflammatory chemokines, such as CCL5 and CXCL10, in tumor cells that promoted the recruitment of T cells into the tumor (*13*). Importantly, there was evidence of immune activation by CDK4/6i in clinical samples from neoadjuvant trials of CDK4/6i abemaciclib, such as neoMONARCH and NeoPalAna (*14, 15*). Finally, several preclinical studies reported that CDK4/6i enhance the efficacy of immune checkpoint blockade targeting PD-1/PD-L1 in mouse models (*16–19*). Despite these promising reports of pro-immunogenic activity, clinical trials combining CDK4/6i with adaptive immune checkpoint inhibitors (e.g., anti-PD-1/PD-L1) have yielded disappointing results, often marked by limited efficacy and increased toxicity (*20*). This failure may be explained by the low mutational burden and limited neoantigen presentation typical of ER+ breast cancers (*21–24*), as well as our previous finding that CDK4/6i can paradoxically reduce the dendritic cell populations necessary for T cell activation (*25*).

Given the limitations of T-cell-based approaches in “cold” breast tumors, we propose shifting the focus toward the innate immune system, specifically Natural Killer (NK) cells, which are potent innate immune effectors. NK cells can recognize stressed or malignant cells independently of MHC-restricted neoantigens. In addition, NK cells are an important source of FLT3L, which can recruit and sustain cross-presenting DCs in the TME, thus initiating adaptive immune responses (*26*). While endogenous pools of NK cells in tumors are limited, they can be expanded *ex vivo* and delivered to patients in the form of cell therapy (*27*). Cell therapy with NK cells is a promising emerging cancer treatment. They provide several advantages over T cell therapies, including a lower risk of cytokine release syndrome (CRS) and graft-versus-host disease (GVHD), and the potential for “off-the-shelf” use from healthy donors (*28–31*).

The success of NK cell therapies in solid tumors is currently hampered by the loss of NK cell activity in the TME. Therefore, a strategy to “prime” the tumor for NK cell recognition is required (*32*). NK cell-mediated recognition of target cells is governed by a balance of signals. Activating receptors, such as NKG2D, recognize stress-induced ligands (e.g., MICA/B and ULBPs) on target cells, while inhibitory receptors (e.g., KIR, NKG2A) recognize MHC-I molecules to prevent autoimmunity (*33–35*). Certain HLAs could also activate NK cells through activating KIRs and NKG2C (*36*). Upregulation of stress ligands on the surface of cells under duress from various stressors, such as viral infection or malignant transformation, serves as a primary “kill” signal, initiating the cytotoxic response of the NK cells (*37, 38*). Another factor that plays a crucial role in NK cell recognition of target cells is intercellular adhesion molecule-1 (ICAM-1). It acts as a critical adhesion molecule on the target cell, enabling the NK cell’s LFA-1 receptor to form a stable and effective connection (*39, 40*). ICAM-1 is an inducible adhesion molecule that can be upregulated by various factors, including cellular stress and inflammation. It is a known downstream target of signaling pathways activated by inflammatory cytokines such as IFN-γ, IFN-α, and TNF-α, which play a role in antiviral responses (*41, 42*). Of note, CDK4/6i are reported to induce expression of endogenous retroviruses (*43*), suggesting that they may upregulate the IFN-ICAM1 axis. We also know that NK cells play a pivotal role in the surveillance of senescent cells (*44*). Finally, a study in a pancreatic cancer model revealed that senescence induced by a combination of CDK4/6i and MEK inhibitors can specifically trigger NK cell-mediated tumor clearance through the production of a SASP-driven secretome (*45*).

Here, we hypothesized that drug-induced stress and senescence caused by CDK4/6i would upregulate stress ligands on tumor cells, which could be leveraged to sensitize ER+ breast cancer to NK cell-mediated killing. To determine whether abemaciclib could transform ER+ breast cancer tumors into viable targets for adoptive NK cell therapy, we investigated the molecular mechanisms by which CDK4/6 inhibition enhances NK cell infiltration and cytotoxicity across a diverse set of ER+ breast cancer patient-derived organoids (PDOs). PDOs have been established as powerful preclinical models for predicting therapeutic response due to their ability to faithfully retain the genetic, phenotypic, and drug sensitivity characteristics of original tumors (*46–57*). We also used patient-derived xenografts (PDX) established in human IL-15-expressing mice to ensure the physiological function of human NK cells in mice, enabling us to test the therapeutic efficacy of CDK4/6i and NK cell combinations *in vivo*.

## RESULTS

### CDK4/6i treatment sensitized breast cancer cells to NK cell-mediated killing

To investigate whether CDK4/6i promotes tumor recognition by NK cells, we assessed the surface expression of a panel of NK cell-activating and inhibitory ligands on ER+ breast cancer cell lines (MCF7, T47D) and patient-derived cells (PDC8798) following treatment with abemaciclib or vehicle. Our analysis revealed that CDK4/6 inhibition significantly upregulated the stress ligands ULBP2/5/6 and MICA/B (Fig. 1A-C), which are known to trigger NK cell cytotoxicity via the NKG2D receptor. Furthermore, we observed a robust increase in the surface expression of the adhesion molecule ICAM-1, a critical factor for the formation of the immunological synapse between NK cells and their targets (Fig. 1A-C).

**Fig. 1.**
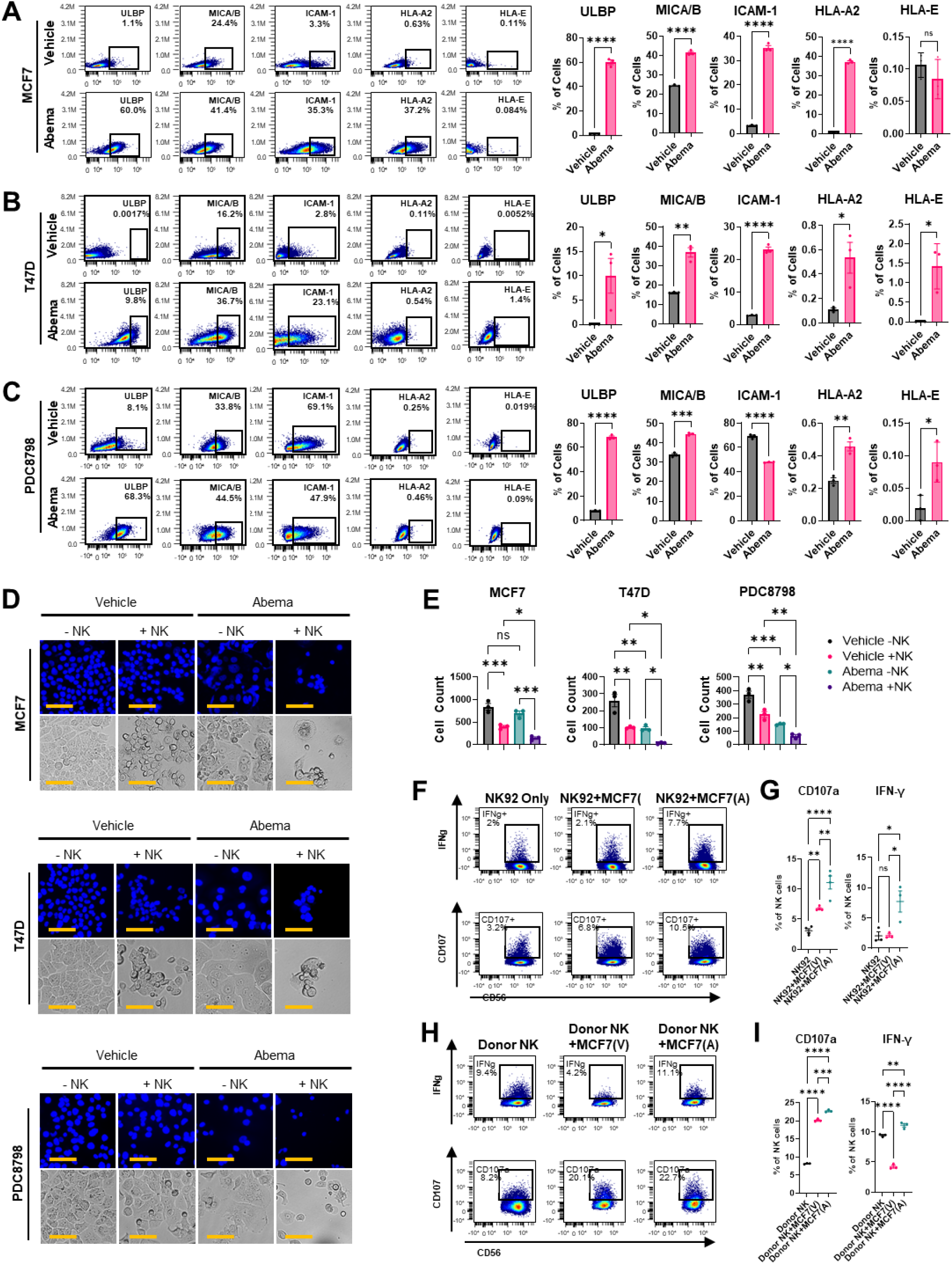
CDK4/6i-treated tumor cells promote NK cell activation. (**A** to **C**) Flow cytometry analysis of cell surface expression of indicated molecules in MCF7 cells (A), T47D cells (B), and PDC8798 (C) treated with vehicle or 1 μM abemaciclib for 5 days. Left panels show representative cytometry plots, and graphs on the right show quantified data. Error bars indicate mean ± SEM. n=3. *p < 0.05; **p < 0.01; ***p < 0.001; ****p < 0.0001(two-tailed t-test). (**D**) Representative microphotographs of MCF7, T47D, and PDC8798 tumor cells after co-culture with NK92 cells. Cells were visualized with DNA dye Hoechst (top panels) and by bright-field microscopy (bottom panels). Cells were pre-treated with vehicle or 1 μM abemaciclib for 5 days and incubated with NK cells for 24 hours. NK cells were removed by washing before imaging. Scale bar: 75μm. (**E**) Quantification of images in (D). Data presented as mean ± SEM. Data are the average of three biological replicates (n = 3). *p < 0.05; **p < 0.01; ***p < 0.001 (one-way ANOVA). (**F**) Flow cytometry analysis of indicated markers in NK92 cells after co-culture with MCF7 tumor cells. MCF7 cells were pre-treated with vehicle or 1 μM abemaciclib for 5 days, then NK92 cells were added for co-culture for an additional 24 hours. (**G**) Quantification of data shown in (F). Error bars indicate mean ± SEM. n=3. *p < 0.05; **p < 0.01; ****p < 0.0001 (one-way ANOVA). (**H**) Same as F, except donor NK cells were used. (**I**) Quantification of data in (H). Data presented as mean ± SEM. n=3. **p < 0.01; ***p < 0.001; ****p < 0.0001 (one-way ANOVA).

Concurrently, we examined the expression of inhibitory MHC class I molecules, specifically HLA-A2 and HLA-E, which can traditionally suppress NK cell-mediated killing. While abemaciclib treatment increased MHC class I expression on MCF7 cells, this effect was not observed in T47D or PDC8798 cells (Fig. 1A-C). Additionally, the expression of HLA-E, the cognate ligand for the inhibitory NKG2A receptor, was not significantly altered by CDK4/6i treatment (Fig. 1A-C). We next performed co-culture assays to determine whether CDK4/6i pre-treatment enhances NK cell-mediated killing of tumor cells. Human ER+ breast cancer cell lines (MCF7, T47D) and patient-derived cells (PDC8798) were pre-treated for 5 days with vehicle or abemaciclib, and then co-cultured with human NK cells (NK92) for an additional 24 hours. We observed a notable reduction in tumor cell numbers after co-culture with NK cells compared with no-NK-cell controls, indicating NK cell-mediated killing (Fig. 1D, E). Importantly, consistent with the induction of NK-activating signals, CDK4/6i-treated cells were more effectively eliminated by NK cells than vehicle-treated cells (Fig. 1D, E).

To determine if the observed ligand upregulation translates to enhanced effector function, we evaluated NK cell activation following co-culture with abemaciclib-treated MCF7 cells. Two types of NK cells were tested: the NK92 cell line and donor-derived *ex vivo* expanded NK cells. We quantified the expression of the inflammatory cytokine IFNγ and the degranulation marker CD107a using flow cytometry. Our analysis revealed that incubation with abemaciclib-pretreated MCF7 cells significantly stimulated both IFNγ production and CD107a translocation in NK92 cells (Fig. 1F, G) and in primary donor NK cells (Fig. 1H, I). These findings demonstrate that the CDK4/6i-induced surface profile triggers NK cell effector programs.

### Establishment of an ER+ BC PDO platform for pre-clinical testing of CDK4/6i and NK cell therapy

To evaluate whether CDK4/6i-induced stress sensitizes breast cancer cells in a clinically relevant context, we established a biobank of patient-derived organoids (PDOs) derived from human breast tumor specimens. We validated the ER status of these models using immunofluorescence (IF) and Western blot analysis (Fig. 2A-C). While all PDOs exhibited ERα expression, we observed significant inter-patient heterogeneity in the percentage of ER-positive cells (Fig. 2A, B). This variation is consistent with clinical observations, where ER-positivity is defined across a broad spectrum (1–100%) in patients diagnosed with hormone receptor-positive disease (*58*). These findings indicate that our PDO biobank recapitulates the histological diversity of the clinical patient population.

**Fig. 2.**
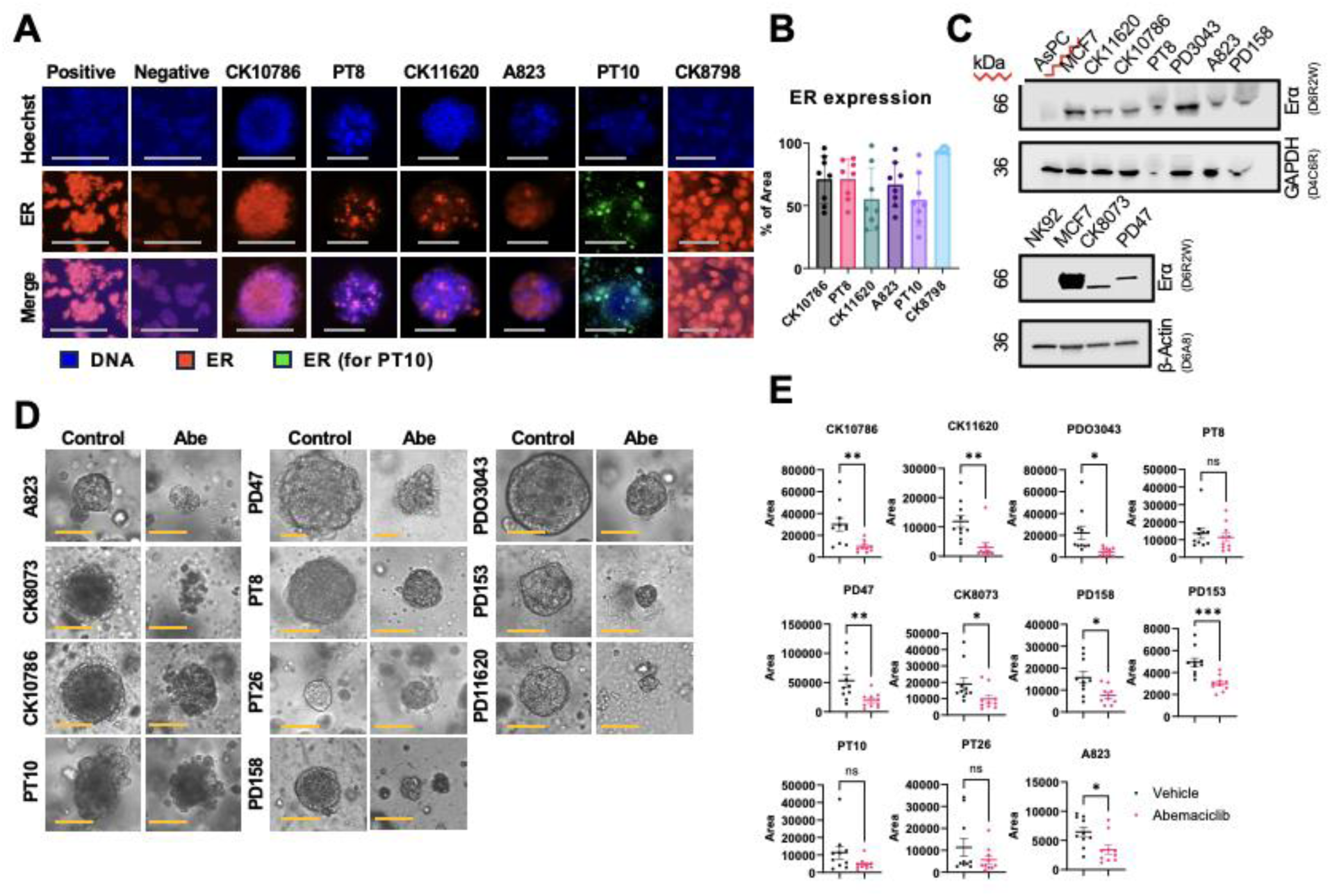
CDK4/6i treatment promotes ICAM-1 and stress ligand expression in PDOs. (**A**) Representative immunofluorescence images of ERα expression in organoids from 5 distinct patients and a patient-derived cell model. Scale bar: 75 μm. (**B**) Quantification of ERα-positive area (%) per organoid in (A). n=8. Data represent mean ± SD. (**C**) Western blot analysis of ERα expression in NK92 cells (negative control), AsPC pancreatic cancer cells (negative control), MCF7 cells (positive control), and PDOs. GAPDH and β-actin were used as loading controls. (**D**) Representative images of PDOs and MCF7 treated with 1 μM abemaciclib or vehicle for 5 days. Images were obtained using transmitted light microscopy on day 5. Scale bar: 75 μm. (**E**) Quantification of PDO area in (D). Data represent mean ± SEM. n=10. *p < 0.05, **p < 0.01 (two-tailed t-test).

We next evaluated the growth inhibitory effects of CDK4/6i across our PDO biobank. Organoids were treated with abemaciclib or vehicle control for five days and monitored via longitudinal live-cell imaging (Fig. 2D). Morphometric analysis of at least 15 individual organoids per group revealed that abemaciclib treatment significantly reduced the total organoid area compared to vehicle controls (Fig. 2E). While the magnitude of growth inhibition varied across the biobank, reflecting the expected diversity in drug sensitivity, a statistically significant reduction in organoid size was observed in 73% (8 of 11) of the tested models. These data confirm that the majority of our patient-derived models were sensitive to the cytostatic effects of abemaciclib, providing a robust platform for subsequent preclinical combination studies with NK cells.

### CDK4/6i promoted the recruitment of NK cells into ER+ breast cancer PDOs and increased tumor cell killing

To determine if CDK4/6i promoted tumor cell clearance by the NK cells, we performed live-cell co-culture assays using pre-labeled ER+ PDOs (CellTracker Green) and NK92 cells (CellTracker Deep Red). Following a 5-day pretreatment with abemaciclib or vehicle, organoids were transitioned to drug-free media and co-cultured with NK cells for 72 hours. We employed longitudinal fluorescence imaging to monitor NK cell infiltration and tumor cell viability (DAPI) at 18, 48, and 72 hours (Fig. 3A, B).

**Fig. 3.**
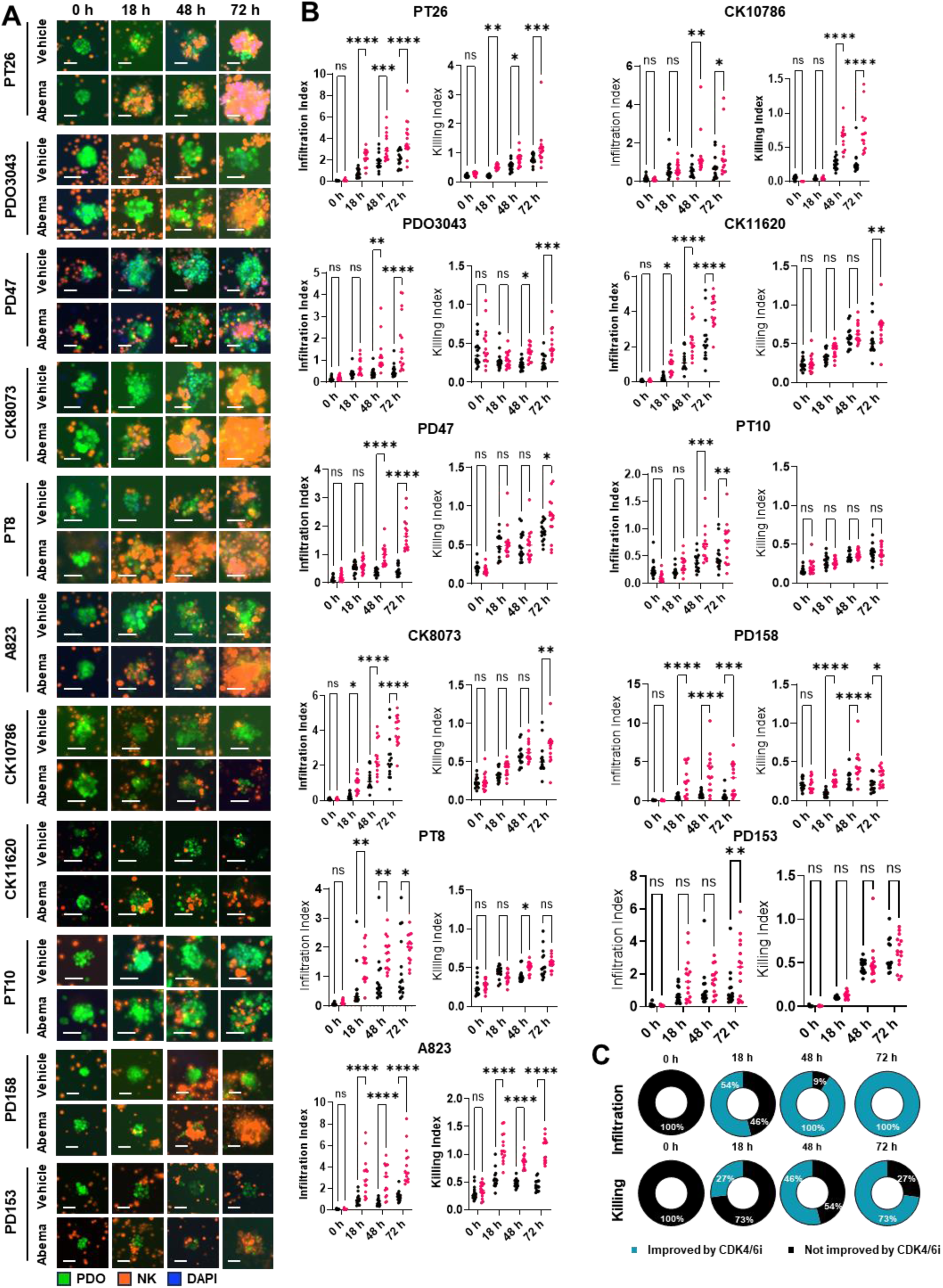
CDK4/6i treatment enhances the interaction between PDOs and NK cells. (**A**) Representative images of breast cancer organoids pre-treated with either vehicle or 1 µM abemaciclib for 5 days, followed by co-culture with NK92 cells. Organoids were labeled with CellTracker Green (green signal), and NK cells were labeled with CellTracker Deep Red (orange). Dead cells are stained with DAPI (blue). Images were acquired at 0, 18, 48, and 72 h. Scale bar: 55 μm. (**B**) Quantification of NK-PDO interactions from (A) using ImageJ. Infiltration index was calculated as the ratio of red-to-green fluorescence (NK cells within PDOs); killing index was defined as blue-to-green fluorescence (dead cells within PDOs). n=15. *p < 0.05, **p < 0.01, ***p < 0.001, ****p < 0.0001 (two-way ANOVA). (**C**) Pie chart summarizing data in (B) to show the proportion of PDOs (n=11 patients) that displayed significantly enhanced NK cell infiltration and killing (blue) versus those that did not (black) across four timepoints.

Quantitative image analysis revealed that abemaciclib-treated PDOs exhibited significantly enhanced NK cell recruitment and accelerated tumor cell death compared to vehicle controls. Notably, increased NK cell infiltration was observed across all tested ER+ PDO models, with 73% (8 of 11) demonstrating a significant increase in NK-mediated killing by the 72-hour endpoint (Fig. 3C). In several models, this potentiation was evident as early as 18 hours post-co-culture, suggesting a rapid recognition of the “primed” tumor state.

### CDK4/6i induced NK cell-interacting cell surface proteins in PDO cells

To dissect the molecular shifts induced by CDK4/6i at single-cell resolution, we dissociated the PDOs and performed spectral flow cytometry for a panel of NK cell-modulating signals. This analysis was performed for all 11 PDO models and representative flow plots from one model are shown in Fig. 4A. In the basal (vehicle-treated) state, the majority of PDO models exhibited minimal surface expression of stress ligands, with a median of 6.2% for ULBPs and 4.6% for MICA/B (Fig. 4B). Notable exceptions included CK10786 and PDO3043, which displayed high baseline levels of ULBPs (58%) and MICA/B (38%), respectively. Following abemaciclib treatment, we observed a robust induction of ULBPs, with 6 of 11 models reaching >50% positivity (Fig. 4B). While the induction of MICA/B was less pronounced than that of ULBPs, 10 out of 11 models showed significant upregulation of at least one activating ligand or the adhesion molecule ICAM-1 (Fig. 4C). Basal ICAM-1 expression was low-to-moderate across the biobank (median 37.6%) but was consistently induced by CDK4/6i treatment (median 44.6%; Fig. 4B).

**Fig. 4.**
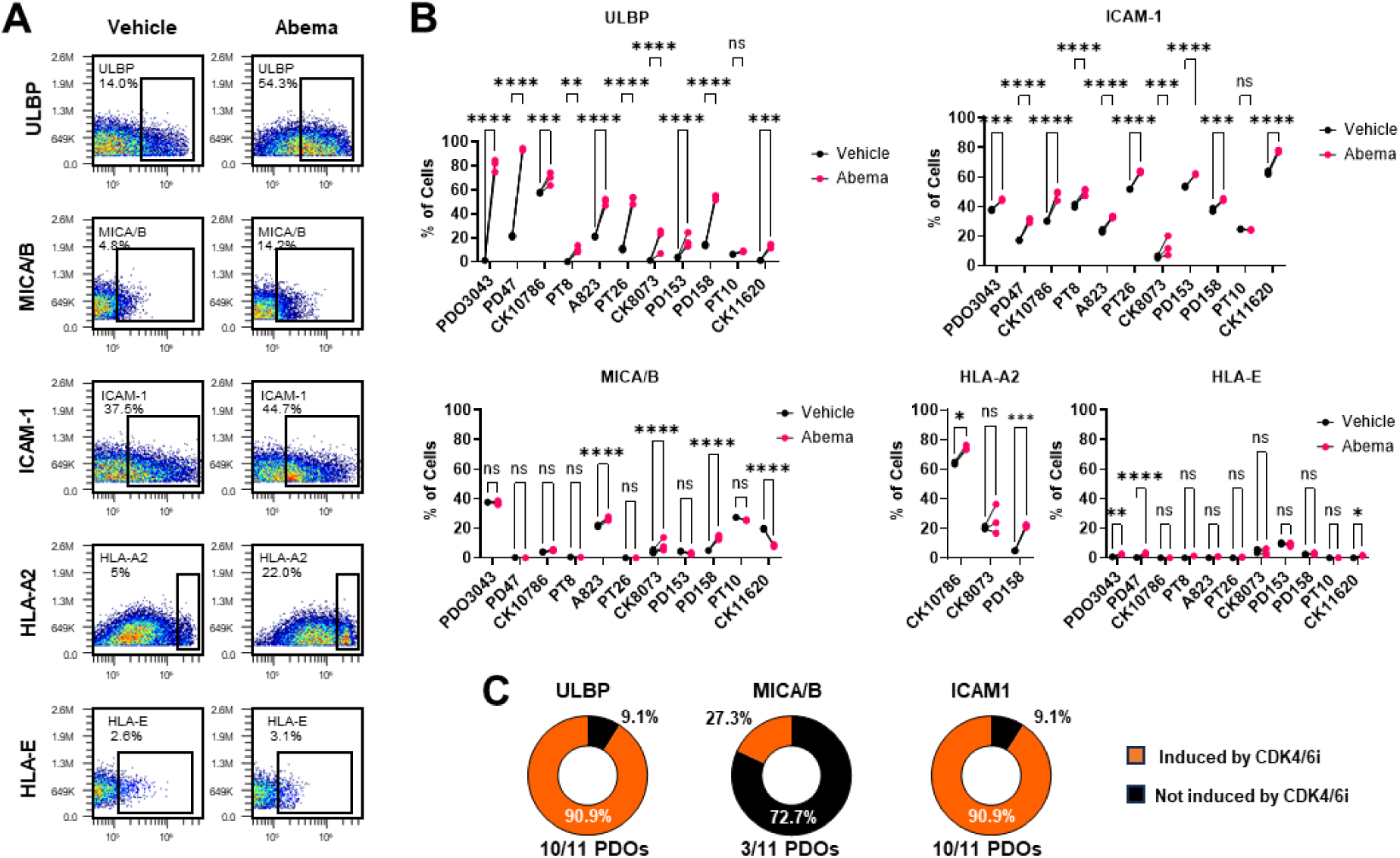
CDK4/6i induced-stress ligands and ICAM-1 on the surface of cancer cells from ER+ PDOs. (**A**) Representative flow cytometry plots showing surface expression of indicated NK cell-activating and inhibiting signals in a PDO158 treated with 1 μM abemaciclib or vehicle control for 5 days. (**B**) Quantification of surface marker expression across 11 PDOs treated as in (A). n=3. ***p < 0.001, ****p < 0.0001 (two-way ANOVA). (**C**) Pie chart showing the proportion of PDOs that upregulated stress ligands and ICAM-1 after CDK4/6i treatment based on data in (B).

We further evaluated the potential for CDK4/6i to induce inhibitory signals. Two of three HLA-A2+ PDOs showed a slight increase in expression (Fig. 4B). The expression of the inhibitory ligand HLA-E remained consistently low and was not significantly altered by treatment (Fig. 4B). Collectively, these findings demonstrate that CDK4/6i shifts the tumor surface profile toward a pro-immunogenic state in the majority of patient models.

### Stress ligands and ICAM-1 were required for NK cell killing of CDK4/6i-treated breast cancer cells

To investigate the functional necessity of abemaciclib-induced surface markers in enhancing NK cell activity, we performed a series of loss-of-function assays using neutralizing antibodies and blocking proteins. MCF7 cells were pretreated with abemaciclib or vehicle, followed by co-culture with NK cells in the presence or absence of two distinct ICAM-1-targeting antibodies. In the abemaciclib-pretreated group, we observed that ICAM-1 blockade significantly inhibited NK-mediated cytotoxicity, whereas no such inhibitory effects were observed in the vehicle-treated groups (Fig. 5A, B). We also used an antibody targeting the NKG2D receptor on NK cells, or an NKG2D recombinant protein that blocks the stress ligands ULBP and MICA/B on tumor cells. Similarly to ICAM-1, blocking either the stress ligands or their cognate receptor, NKG2D, significantly impaired NK-mediated tumor cell killing (Fig. 5A, B). These data suggest that ICAM-1 and stress ligands are important for CDK4/6i-mediated NK cell cytotoxicity towards tumor cells.

**Fig. 5.**
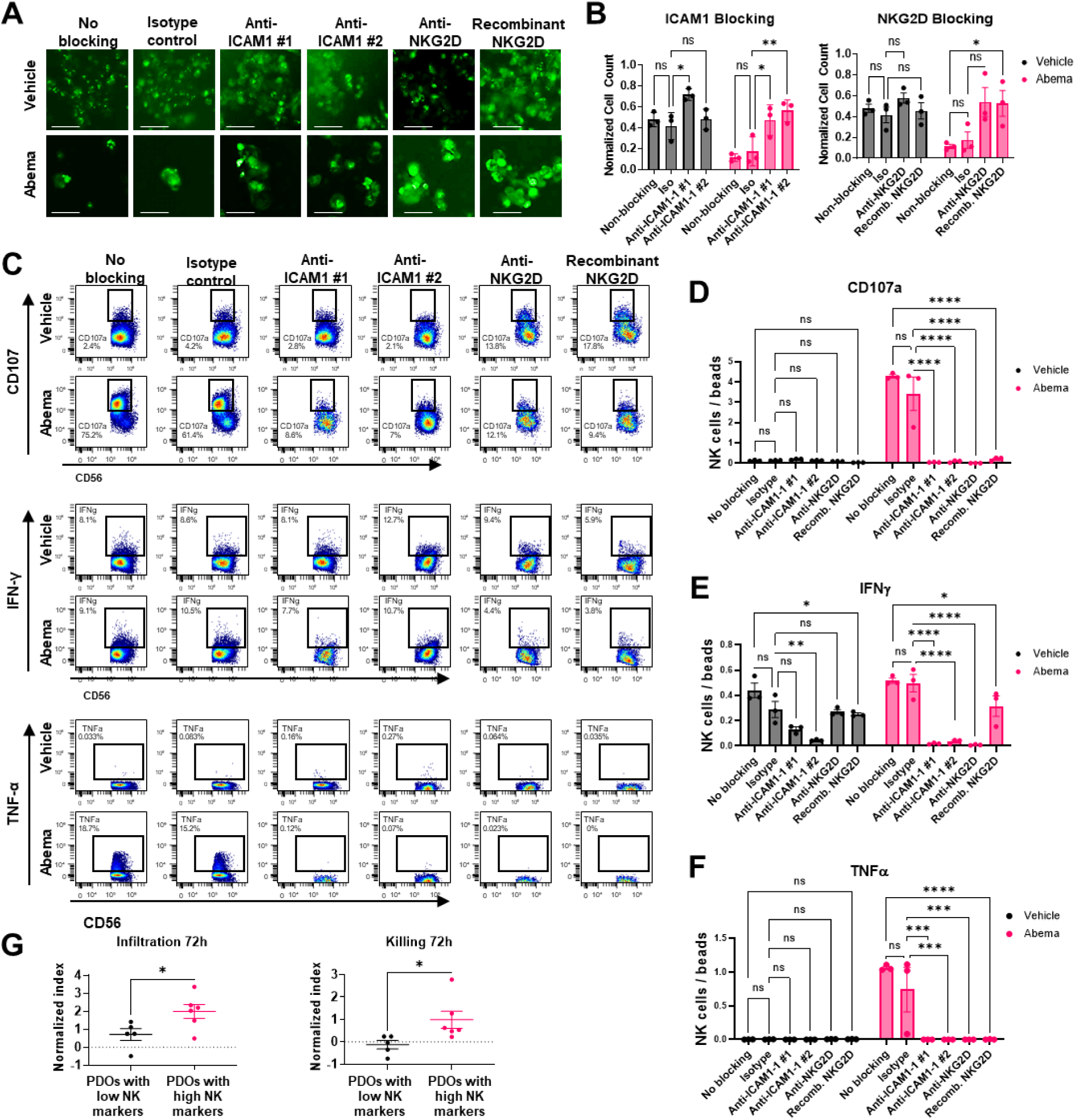
CDK4/6i-induced stress ligands and ICAM-1 are required for NK cytotoxicity towards tumor cells. (**A**) Representative images of 5-day vehicle or 1 μM abemaciclib-pretreated MCF7 cells after co-culture with NK cells in the absence or presence of indicated blocking reagents. Live cells were stained with Calcein AM (green). Scale bar: 55 μm. (**B**) Quantification of co-culture assays in (A). Cell viability was normalized to tumor-only controls (vehicle or abemaciclib pretreatment) to specifically assess NK-mediated effects. Error bars indicate mean ± SD. n=3. *p < 0.05; **p < 0.01 (one-way ANOVA). (**C**) Representative flow cytometry plots showing surface expression of indicated markers in NK92 cells after 24 h co-culture with MCF7 cells pretreated with vehicle or abemaciclib (1 μM) for 5 days. Blocking reagents were added as indicated. (**D, E, F**) Quantification of flow cytometry data shown in (C). Cell counts were normalized to cell counting beads. Error bars indicate mean ± SD. n=3. *p < 0.05; **p < 0.01 (one-way ANOVA). (**G**) Correlation between ICAM-1/ULBP expression levels (low vs. high) and NK infiltration and killing index at 72 h in PDO-NK co-cultures (from Fig. 3). Error bars indicate mean ± SD. *P < 0.05 two-tailed t-test.

Beyond direct cytotoxicity, we evaluated whether interactions with ICAM1 and stress ligands on the tumor cell surface were required for the induction of NK cell effector phenotype. Flow cytometry profiling revealed that NK cells co-cultured with abemaciclib-primed tumor cells in the presence of ULBP or ICAM-1 blockers exhibited significantly reduced expression of the degranulation marker CD107a and immune response-promoting cytokines IFN-γ and TNF-α (Fig. 5C-F). This shows that CDK4/6i-induced immune-interacting surface ligands are important for activating NK cell effector function.

To determine whether the requirement for abemaciclib-induced stress ligands and ICAM-1 in NK cell recruitment and cytotoxicity is consistent across PDO models, we performed a correlation analysis across our PDO biobank. We found that PDOs characterized by high surface levels of ULBP and ICAM-1 exhibited significantly greater NK cell infiltration and tumor cell killing at the 72-hour co-culture time point compared to low-expressing models (Fig. 5G).Together, these results identify ICAM-1 and NKG2D ligands as key CDK4/6i-induced sensitizers that are functionally required for NK cell-mediated cytotoxicity.

### CDK4/6i induced stress ligands and ICAM-1 via divergent signaling pathways

We next explored the molecular mechanisms underlying the CDK4/6i-mediated induction of stress ligands and adhesion molecules. Based on our previous findings that PI3K/mTOR axis fueled the inflammatory signaling in CDK4/6i-treated cells (*8*), and the established role of immune transcription factors NF-κB and STAT1 in stress response and SASP, we performed a loss-of-function experiment using inhibitors of these factors. MCF7 cells were treated with abemaciclib alone or in combination with BMS-345541 (NF-κB inhibitor), gedatolisib (dual PI3K/mTOR inhibitor), or ruxolitinib (JAK/STAT1 inhibitor) for five days. Our analysis revealed a clear mechanistic divergence in the CDK4/6i-mediated regulation of surface immune-interacting proteins. The induction of ICAM-1 was primarily dependent on inflammatory signaling as treatment with the NF-κB inhibitor significantly abrogated ICAM-1 surface expression, as well as, total protein and transcript levels (Fig. 6A, B, C). While STAT1 inhibition also reduced ICAM-1 surface and total protein levels, it did not significantly impact its mRNA expression, suggesting a post-transcriptional regulatory component (Fig. 6A, B, C).

**Fig. 6.**
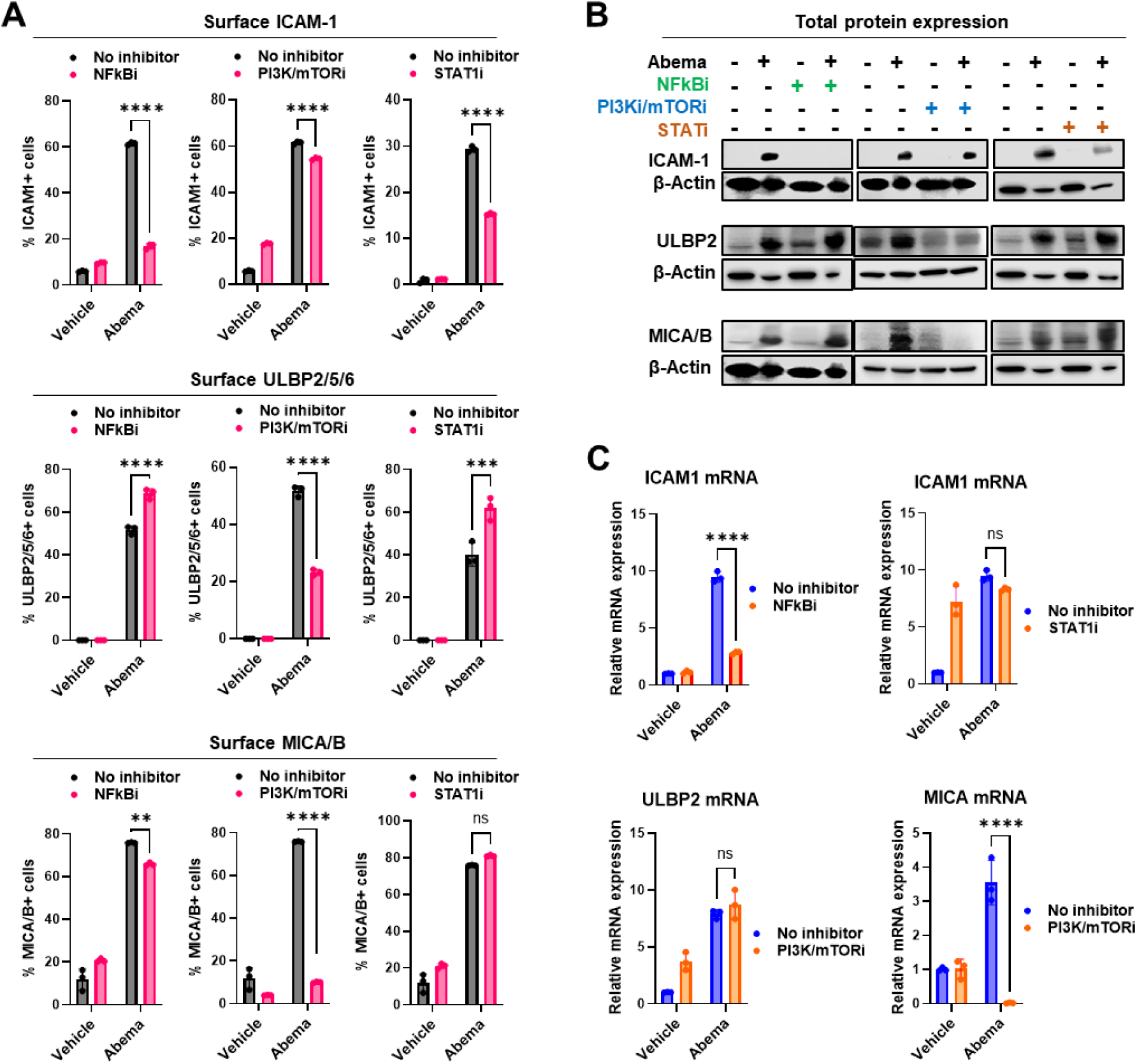
CDK4/6i-induces stress ligands and ICAM-1 via divergent molecular pathways. (**A**) Flow cytometric analysis of surface expression of ICAM-1 (top), ULBP2/5/6 (middle), and MICA/B (bottom). MCF7 cells were pretreated with vehicle, abemaciclib, or the indicated inhibitors, or with combinations thereof, for 5 days. Quantification of the indicated molecules’ expression was shown adjacent to the plots. Error bars indicate mean ± SD. n=3. *P < 0.05; **P < 0.01 (two-way ANOVA). Statistical analysis was performed using two-way ANOVA; data represent mean ± SD, ***p < 0.001, ****p < 0.0001. (**B**) Western blot analysis of ICAM-1, ULBP2, and MICA/B protein expression in MCF7 cells treated with vehicle, abemaciclib, NF-κB inhibitor, or their combination. β-actin was used as a loading control. (**C**) Relative ICAM-1, ULBP2, and MICA mRNA expression quantified by qPCR. Statistical analysis was performed using two-way ANOVA; data represent mean ± SD, ***p < 0.001, ****p < 0.0001.

Conversely, the induction of stress ligands was more closely tied to metabolic signaling. The upregulation of surface ULBP2/5/6 and MICA/B was significantly attenuated by dual PI3K/mTOR inhibition (Fig. 6). Interestingly, while both MICA protein and transcription were PI3K/mTOR-dependent, ULBP2 protein levels were reduced without a concomitant decrease in mRNA, suggesting that PI3K/mTOR signaling may regulate certain stress ligands via post-transcriptional mechanisms. We also observed that NF-κB inhibition modulated surface stress ligand expression. However, these changes were modest and the protein levels were minimally affected (Fig. 6). Collectively, these results demonstrate that CDK4/6i-mediated NK sensitization is driven by a bifurcated signaling network: NF-κB signaling primarily orchestrates the induction of the adhesion molecule ICAM-1, while the PI3K/mTOR pathway is the predominant regulator of stress ligand expression.

### NK-associated markers display favorable clinical correlations in ER+ breast cancer

To assess the clinical relevance of our findings, we examined the expression of ICAM-1 and NKG2D ligands in human breast cancer specimens using the Human Protein Atlas database. Consistent with our untreated PDO results, immunohistochemical data revealed that ICAM-1 expression is largely low (72.7%) or moderate in primary tumors, while ULBP1 and ULBP3 were undetectable in over 90% of samples (Fig. 7A-D). Furthermore, the NK-activating receptor NKG2D was not detectable within the tumor microenvironment (TME) of most samples (Fig. 7C), suggesting a baseline “cold” innate immune landscape.

**Fig. 7.**
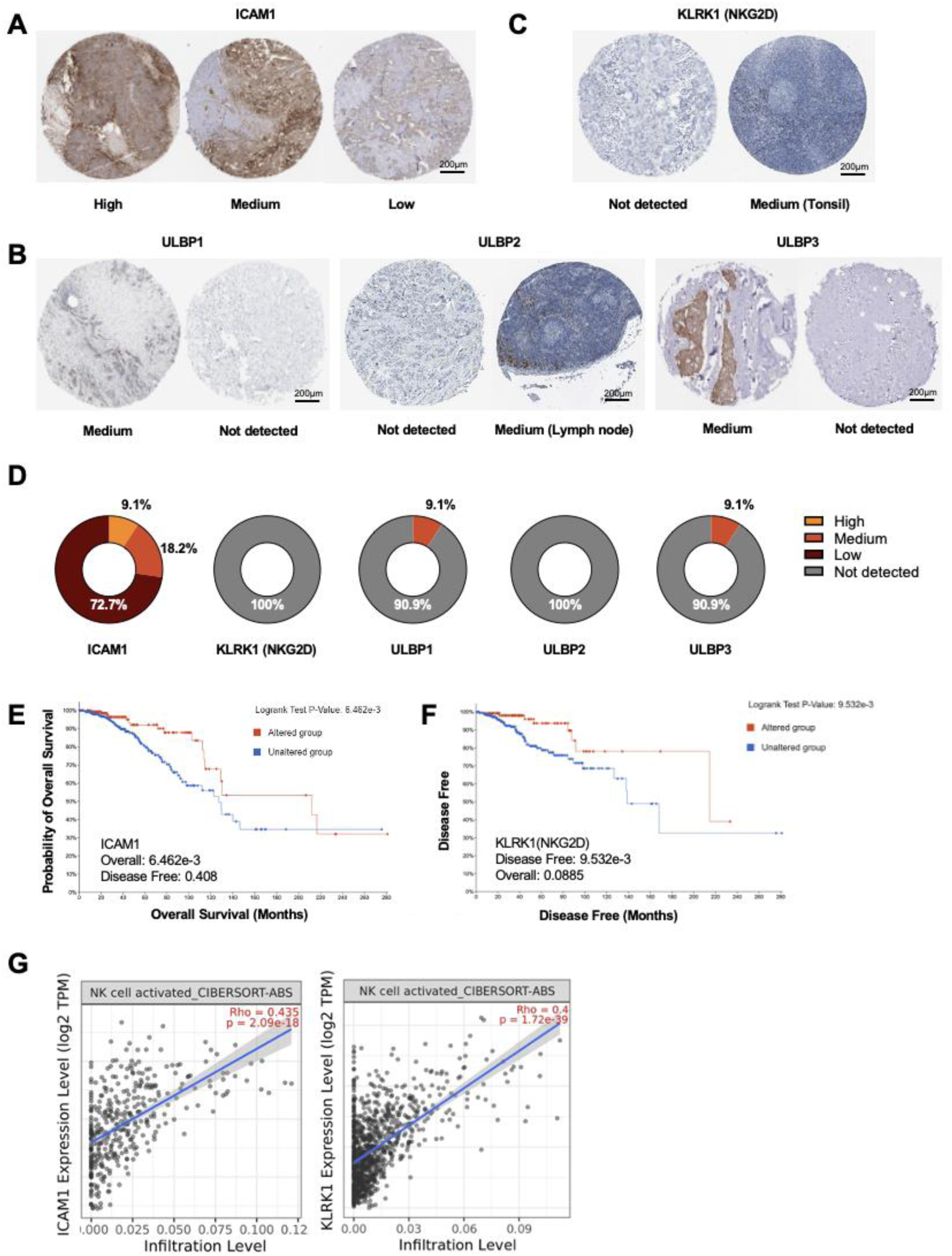
Higher ICAM-1 and NKG2D expression is associated with better prognosis in human breast cancer. (**A**, **B**, **C)** Representative immunohistochemistry staining patterns of ICAM1, NKG2D (KLRK1), ULBP1, ULBP2, and ULBP3 in breast cancer specimens, with tonsil and lymph node tissues serving as positive controls, obtained from the Human Protein Atlas (HPA, http://www.proteinatlas.org/). Examples of high, medium, and low/undetectable expression are shown. Scale bar: 200 μm. (**D**) Pie chart showing the distribution of protein expression levels from the immunohistochemistry results in (A). (**E**) Breast cancer patients with high ICAM1 expression exhibited improved overall survival (p < 0.01) compared with patients with low ICAM1 expression, as determined by the log-rank test using data from cBioPortal for Cancer Genomics. (**F)** Breast cancer patients with high KLRK1 (NKG2D) expression showed longer disease-free survival (p < 0.01), as analyzed by the log-rank test via cBioPortal for Cancer Genomics. (**G**) Scatter plots showing the correlation between NK cell infiltration estimates and ICAM1 (left) or KLRK1 (right) gene expression in breast cancer, as calculated by TIMER2.0 using the CIBERSORT-ABS algorithm and Spearman correlation analysis.

Using cBioPortal and TCGA data, we investigated the impact of these factors on patient outcomes. In ER+ breast cancer, high ICAM1 expression was significantly associated with improved overall survival (Fig. 7E), and elevated KLRK1 (NKG2D) expression correlated with longer disease-free survival (Fig. 7F). Furthermore, TIMER 2.0 analysis confirmed that both ICAM1 and KLRK1 levels positively correlate with NK cell infiltration and activation markers (CIBERSORT-ABS; Fig. 7G). These clinical data underscore that the NK cell activating factors upregulated by CDK4/6i in our models, specifically ICAM-1 and NKG2D ligands, are associated with NK cell infiltration and superior clinical outcomes in patients.

### CDK4/6i enhances the efficacy of NK cell therapy *in vivo*

We next evaluated the efficacy of combining a CDK4/6 inhibitor and NK cell therapy *in vivo*. We used female NSG-SGM-huIL15 mice, which provide physiological levels of human IL15 (7.1 +/- 0.3 pg/ml) to support human NK engraftment (*59*). This mouse model enables the study of human NK cells and their efficacy against human tumors *in vivo*. Mice were implanted with a slow-release human estrogen pellet to enable growth of human ER+ breast cancer tumors and treated with CDK4/6i and NK cells in either sequential (Fig. 8A) or concurrent treatment schedule (Fig. 8B).

**Fig. 8.**
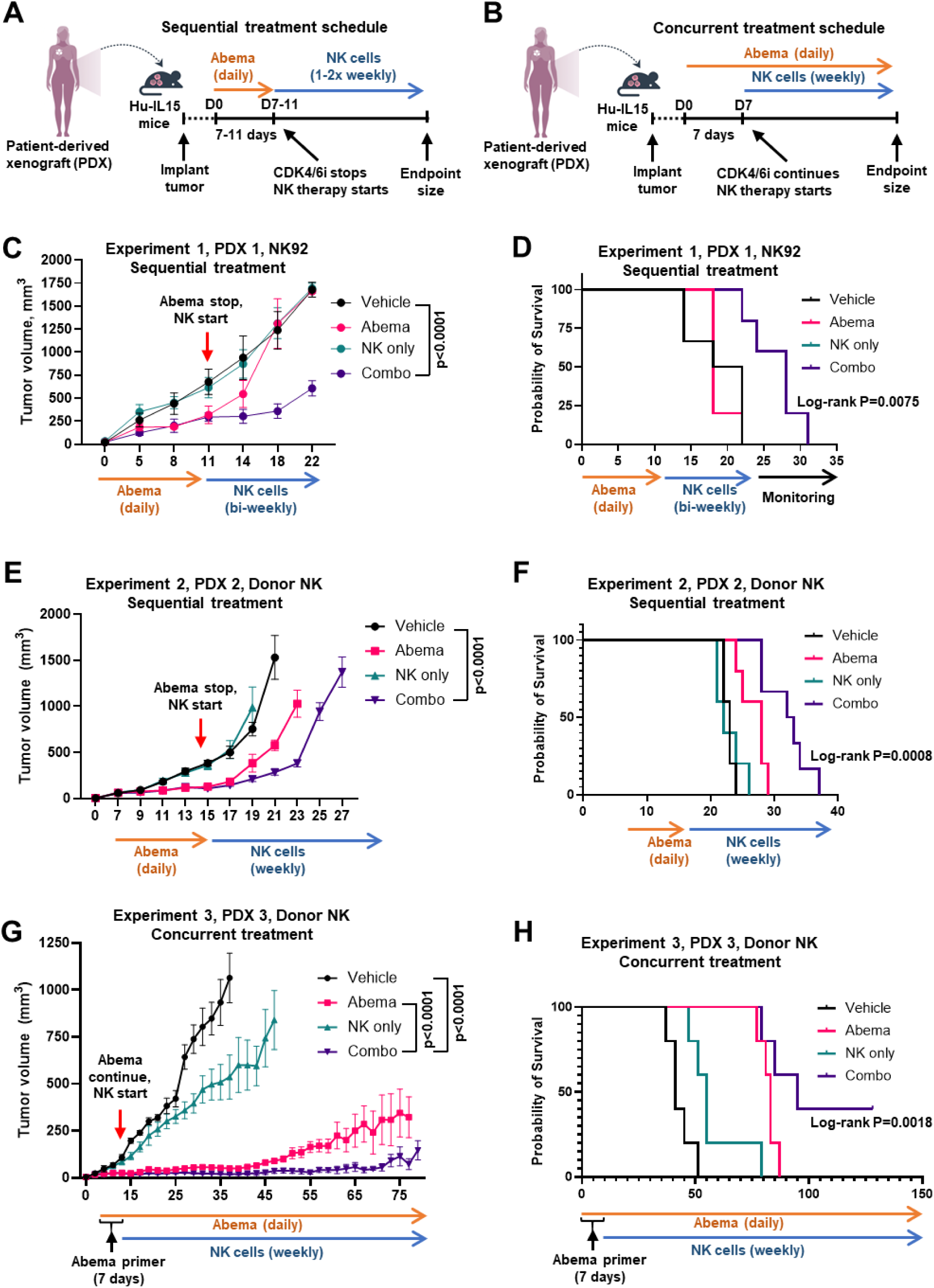
Analysis of the efficacy of NK cell therapy after CDK4/6i treatment *in vivo*. (**A, B**) Schematic of in vivo experiments administering CDK4/6i and NK cells sequentially (A) or concurrently (B). For sequential treatment (experiments 1 and 2, C-F), mice were pretreated with abemaciclib for 7-11 days, followed by a single NK cell infusion (10×10⁶ cells/injection). For concurrent treatment (experiment 3, G-H), mice were pretreated with abemaciclib for one week, and abemaciclib and NK cells (10×10⁶ cells/injection, weekly) were administered simultaneously. (**C**) Experiment 1, sequential treatment. Tumor growth curves in PDX-bearing mice treated with abemaciclib (75 mg/kg, daily) and NK92 cells. The treatment schedule is as shown in (A). n = 5 mice/group. Statistical significance calculated using 2-way ANOVA with Tukey’s post-test. (**D**) Corresponding survival curves for the experiment in (C). The Log-rank (Mantel-Cox) test was used to compare the combo and vehicle groups. (**E-F**) Tumor growth and survival in Experiment 2 with sequential treatment. The treatment scheme is shown in (A). Doses and analyses as in (C-D). n = 5 mice/group. (**G-H**) Tumor growth and survival in Experiment 3 with concurrent treatment. The treatment scheme is shown in (B). Doses and analyses as in (C-D). n = 5 mice/group.

In our initial cohort, we tested a sequential treatment regimen (Fig. 8A): mice were pretreated with abemaciclib for seven days, followed by drug withdrawal and subsequent NK92 cell infusion. We found that NK cell infusions into the untreated tumors had minimal impact. In the abemaciclib-only group, tumor growth was initially inhibited, but tumors rapidly resumed growth upon treatment cessation, eventually reaching volumes comparable to vehicle-treated controls. Notably, combination treatment significantly improved tumor growth inhibition and survival compared to either monotherapy (Fig. 8C-F). We observed a similar trend when utilizing primary donor-derived expanded NK cells in another PDX model. However, we also noted that while survival was significantly prolonged, the sequential regimen did not achieve sustained tumor regression (Fig.8C-F).

We next tested a concurrent treatment strategy, postulating that it would enable tumors to maintain the “primed” stress-ligand-enriched surface phenotype. In this second dosing schedule, mice received a seven-day abemaciclib lead-in followed by donor NK cell infusions while maintaining continuous CDK4/6i therapy (Fig. 8B). Under concurrent conditions, abemaciclib monotherapy initially induced marked growth inhibition. However, tumors eventually escaped this arrest, mirroring the acquired resistance observed in clinical settings (Fig. 8G, 8H). Notably, the addition of NK cells significantly delayed this regrowth compared to abemaciclib alone. Furthermore, a subset of mice treated with CDK4/6i and NK cells achieved complete tumor regression (40%). These results indicate that NK cell therapy not only synergizes with CDK4/6i during the initial response phase but also helps delay the onset of therapeutic resistance.

## DISCUSSION

The main goal of this study was to explore whether a small-molecule CDK4/6 inhibitor can enhance the anti-tumor activity of innate immune effectors, such as NK cells. We demonstrated that CDK4/6i sensitizes human ER-positive breast cancer cells to NK cell-mediated killing by upregulating stress ligands and adhesion molecule ICAM-1. This study provides pre-clinical evidence that NK cell therapies, such as infusions of healthy donor NK cells and, potentially, CAR-NK cells, can be harnessed to eliminate CDK4/6i-treated tumors. Such combinations offer a promising strategy to convert transient tumor stabilization in metastatic patients on CDK4/6i into durable clinical clearance. Furthermore, this innate-focused strategy is suitable for “cold” tumors that are typically resistant to checkpoint blockade, such as ER+ breast cancer (*60, 61*), because it bypasses the requirement for MHC-restricted antigen presentation and leverages NKG2D-mediated recognition of “induced self” stress markers.

We also detected MHC-I upregulation induced by CDK4/6i in some PDO models. Although elevated MHC-I could, in theory, enhance inhibitory signaling in endogenous NK cells through KIR interactions, it is unlikely to compromise the efficacy of allogeneic NK cell therapy. In the allogeneic setting, donor selection can be strategically optimized to ensure a KIR-HLA mismatch, thereby bypassing MHC-I-mediated inhibition or by choosing a donor with the right combination of HLA and activating KIR.

Our investigation into the molecular mechanisms underlying CDK4/6i-induced NK cell-activating cell-surface phenotype revealed a complex regulatory network. We found that ICAM-1 induction is primarily dependent on NF-κB signaling, whereas the upregulation of stress ligands (ULBPs and MICA/B) is closely linked to metabolic signaling through the PI3K/mTOR pathway. This bifurcated control suggests that CDK4/6i leverages two distinct molecular “engines” to re-engineer the tumor surface for NK cell recognition and clearance. This mechanistic divergence suggests that the immune-sensitizing effects of abemaciclib are deeply integrated into the cell’s metabolic and inflammatory stress responses. Notably, the involvement of the mTOR pathway has significant clinical implications. mTOR inhibitors, such as everolimus, are currently used in advanced HR+ breast cancer management (*62*). However, our data suggest that mTOR inhibition may suppress stress ligand expression, potentially due to the global protein synthesis. Consequently, patients on mTOR inhibitors may be suboptimal candidates for NK-based therapy, underscoring the need for careful pharmacological stratification in combination trials.

Our functional studies revealed that CDK4/6i-induced expression of stress ligands and ICAM-1 is critical for NK cell-mediated cytotoxicity. On the other hand, a key finding in our PDO biobank analysis was the baseline heterogeneity of these markers in tumors from different patients. We propose that surface levels of stress ligands and ICAM-1 could serve as predictive biomarkers for patient stratification. For instance, tumors with high baseline ligand expression may be candidates for NK cell monotherapy, while those with low expression could be sensitized via CDK4/6i priming. This precision approach can help reduce unnecessary drug exposure and associated toxicities.

While our PDX models are established in immunodeficient mice, the combination of CDK4/6i and NK cell therapy is likely to have a broad impact on the tumor microenvironment (TME) in the context of a functional immune system. Previous studies have demonstrated that CDK4/6 inhibition enhances activation of tumor-infiltrating T cells and suppresses regulatory T cell proliferation (*19, 43, 66*). Our previous findings showed that CDK4/6 inhibitor-treated breast cancer cells produce T cell-recruiting chemokines (*8*). Coupled with our new results presented here, this suggests that CDK4/6i creates a permissive environment for both innate and adaptive immunity. This indicates potential broader applicability of CDK4/6i-mediated immune sensitization of tumors beyond NK cell therapies, for example, by combining them with autologous T cells and dendritic cell vaccines (*25*), particularly in high-tumor-mutation-burden contexts.

While our findings establish a compelling preclinical basis for the “prime and kill” strategy where CDK4/6i are used to “prime” tumors for NK cell-mediated killing, several limitations should be noted. First, although our use of a diverse PDO biobank and humanized huIL-15 PDX models provides high translational fidelity, these models do not fully recapitulate the complexity of the human tumor microenvironment (TME). In patients, the presence of immunosuppressive myeloid cells and regulatory T cells may pose additional barriers to NK cell infiltration and persistence that are not fully captured in our human IL-15 PDX models. Second, our study focused on the adaptive induction of immune-interactive cell-surface proteins within a specific window of CDK4/6i treatment; however, the long-term effects of chronic dosing on NK cell fitness in humans remain to be clinically validated.

To bridge our preclinical findings to clinical application, several critical steps are required. Phase I/II clinical trials are necessary to determine the optimal timing for “priming” in humans and to monitor for potential “on-target, off-tumor” toxicities, as NKG2D ligands may be expressed in non-malignant tissues under stress. Furthermore, standardizing the manufacturing and scale-up of NK cell products will be essential for widespread implementation. Future efforts should also focus on validating ICAM-1 and ULBP2/5/6 as predictive biomarkers in patient biopsies to prospectively identify those most likely to benefit from this combination.

In summary, our data establish a pre-clinical rationale for combining CDK4/6i with NK cell therapy. By identifying ULBP and ICAM-1 as mechanistic sensitizers and potential biomarkers for patient stratification, we lay the groundwork for clinical trials that may improve therapeutic response and delay progression of metastatic breast cancer.

## MATERIALS AND METHODS

### Cell lines, Patient-Derived cells, and donor NK cells

MCF7, T47D, and NK92 cells were purchased from the American Type Culture Collection. MCF7 cells were cultured in Dulbecco’s modified Eagle’s medium/F12 medium (DMEM/F12) supplemented with 4.5g/L D-Glucose, L-Glutamine, 10% fetal bovine serum (FBS), and 1% penicillin-streptomycin. T47D cells were cultured in RPMI-1640 supplemented with 10% fetal bovine serum (FBS) and 1% penicillin-streptomycin. NK92 cells were cultured in Alpha Modification of Eagle’s Medium (AMEM) supplemented with GlutaMAX, 10% horse serum, 10% fetal bovine serum (FBS), 4mg Folic Acid, 20mg Inositol, Mercaptoethanol, and 1% penicillin-streptomycin. All media from Gibco. Patient-derived cells, such as PDC8798, were obtained from the NCI Patient-Derived Models Repository (PDMR). These cells were cultured in media prepared according to the recipe provided by PDMR. Briefly, cells were cultured in DMEM-F12 (Gibco) media supplemented with 5% FBS (Gibco), Y-27632 (Tocris), adenine (MilliporeSigma), hydrocortisone, and EGF (PeproTech). Human Donor NK cells and mIL21 feeder cells were kindly provided by Dean Lee’s Lab (Nationwide Children’s Hospital, Columbus, Ohio) and kept in culture as described in Jove (DOI: 10.3791/2540-v).

### Patient-Derived Organoids and Patient-Derived Xenografts

PDOs were established from breast cancer patient tissues provided by Surgical Oncology at The Ohio State University. Additional PDOs were purchased from NCI Patient-Derived Models Repository (PDMR). PDOs PT26, PT8, and PT10 were kindly gifted by Dr. Taru Muranen’s lab at Harvard Medical School (Boston, Massachusetts, USA) (*67*).

Organoids were cultured in complete media recommended by the NCI Patient-Derived Models Repository (PDMR). Briefly, organoids were cultured in DMEM-F12 (Gibco) media supplemented with 10% FBS (Gibco), 1X B27-supplement (Gibco), R-spondin, neuregulin, noggin, FGF10, EGF (all from PeproTech), N-acetylcysteine, nicotinamide, hydrocortisone, 17β-estradiol (all from Sigma), Y-27632 (Tocris), and embedded in basement membrane extract (R&D) in ultra-low attachment plates. PDOs PT26, PT8, and PT10 were cultured as described previously (*67*).

### Reagents (drugs and blocking antibodies)

Small molecule inhibitors, including CDK4/6i abemaciclib and ribociclib, PI3K/mTORi gedatolisib, and NFkB inhibitor BMS-345541 were purchased from AdooQ, MedChemExpress, and Selleckchem. For blocking experiments, specific blocking reagents (Biolegend), including anti-MICA/B, anti-ICAM-1, anti-NKG2D antibody, recombinant NKG2D, and an isotype-matched control antibody (mouse IgG1κ from Invitrogen) were used at a final concentration of 20 μg/mL.

### Immunofluorescent staining and Western Blot

Immunofluorescence analysis of the estrogen receptor was performed using organoids that had been released from BME with organoid harvesting solution. The cells and organoids were fixed with cold 4% paraformaldehyde in PBS on ice for 15 minutes, permeabilized with 0.1% Triton™ X-100 for 20 minutes, and blocked with 1% BSA for 1 hour at room temperature. Primary antibodies ERα were applied at a dilution of 1:500 and incubated at 4 °C overnight. Goat anti-rabbit secondary antibodies conjugated with Alexa Fluor 647 or Alexa Fluor 488 were used at a dilution of 1:2000. Nuclei were stained with 1 µg/mL Hoechst 33342 prior to imaging.

For the western blot, cells or PDOs were lysed in RIPA buffer (MilliporeSigma) supplemented with protease inhibitor cocktail (MilliporeSigma). Protein concentrations were determined using the Bradford Protein Assay (Bio-Rad) according to the manufacturer’s instructions. Equal amounts of protein were resolved by SDS-PAGE and transferred onto PVDF membranes. Membranes were blocked and incubated overnight at 4°C with primary antibodies against ERα, ICAM-1, ULBP2, and MICA/B (Cell Signaling) at a dilution of 1:1,000. The housekeeping proteins β-actin and GAPDH (Cell Signaling) were used as loading controls at a dilution of 1:3,000. After washing, membranes were incubated with HRP-linked secondary antibodies (Cell Signaling) at a dilution of 1:10,000 for 2 hours at room temperature.

### NK-tumor co-culture

MCF7, T47D, and PDC8798 cells were pretreated with vehicle or 1 µM abemaciclib for 5 days, then seeded in 96-well plates at a density of 3,000 cells per well in triplicate and incubated overnight to allow attachment. NK92 cells were subsequently added to the cultures, with or without prior treatment, and co-incubated for 24 hours. Tumor-only wells were included as controls. After incubation, all wells were washed 3 times with PBS, and MCF7 cells were labeled with Hoechst 33342 for imaging.

Patient-derived organoids were pretreated for 5 days with either vehicle or 1 µM abemaciclib. PDOs were then released from BME and stained with 7 µM CellTracker Green for 1 hour at 37°C. NK92 cells were similarly stained with 5 µM CellTracker Deep Red for 30 minutes at 37°C. Following staining, PDOs were co-cultured with NK92 cells under drug-free conditions for up to 72 hours. 0.1 µg/mL DAPI was added to detect DNA in dead cells with compromised membrane integrity. EVOS fluorescence microscopy was performed at 0, 18, 48, and 72 hours of co-culture. Fluorescence signals were quantified using ImageJ. Specifically, microphotographs of organoids were obtained in the CellTracker Green, CellTracker Deep Red, and DAPI channels. Organoids were circled based on the green signal, and all fluorescence signals were measured within the circled area.

### qPCR

Total RNA was isolated from MCF7 cells using RNeasy kit (QIAGEN) according to the manufacturer’s instructions. cDNA was synthesized from 1 µg of total RNA using iScript™ cDNA Synthesis Kit from Bio-Rad following the manufacturer’s protocol. Quantitative real-time PCR was performed using SYBR™ Green qPCR Master Mix from Fisher. The primers of GAPDH(F: 5’-CCA TGG AGA AGG CTG GGG-3’, R: 5’-GGT CAT GAG TCC TTC CAC GA-3’), ICAM-1 (F: 5’-GTA TGA ACT GAG CAA TGT GCA AG-3’, R: 5’-GTT CCA CCC GTT CTG GAG TC-3’), ULBP2 (F: 5’-CCT AGC GCT CTG GGT CCTT-3’, R: 5’-AAA GAG AGT GAG GGT CGG CTC-3’), and MICA (F: 5’-GCA GAA ACA TGG AAT GTC TGC CAA-3’, R: 5’-TTC CTA CTT CTG GCT GGC ATC-3’) were purchased from IDT. GAPDH was used as an internal control for normalization. Relative gene expression was calculated using the 2^-ΔΔCt method.

### Flow cytometry

Cells were processed for viability staining with Fixable Viability Dye eFluor 780 according to BD Biosciences’ instructions. Fc receptor blocking was performed with anti-human Fc receptor antibody for 30 minutes, followed by staining with fluorescently labeled antibodies against cell markers. Details of antibody panels, including clones, fluorophores, and vendors, are listed in the resources table. For intracellular staining, cells were fixed and permeabilized using the IC Fixation Buffer and Permeabilization Buffer from Thermofisher. After staining, cells were resuspended in 0.5% buffered formaldehyde and analyzed on a 5-laser Cytek Aurora (Cytek Biosciences) for tumor ligand expression or NK cell activity markers.

### Mouse experiments

All animal experiments were approved by the Ohio State University IACUC (protocol# 2022A00000003). NSG-Tg(Hu-IL15) mice were purchased from The Jackson Laboratory (JAX stock #030890) and bred at the OSU ULAR facility. Female mice were used in this study as breast cancer primarily affects the female population. For the transplanted model, mice were 7-10 weeks of age at the time of tumor inoculation. Patient-derived xenografts (PDX) were cut into 2 mm^3^ fragments and implanted into the 4th mammary fat pad on female hu-IL15 mice, and 17β-estradiol pellets were implanted subcutaneously. Mice will be randomized for therapy with abemaciclib or vehicle. Abemaciclib was administered by daily oral gavage at 75 mg/kg. The oral solution was prepared in 0.5% hydroxyethylcellulose. Infusions of NK cells were delivered once a week at 10 × 10^6 cells via a retroorbital i.v. injection. Tumors were measured using calipers 2-3 times per week, and tumor volume was calculated as 0.5 × length × width × width (V = (L * W^2^)/2). Mice were euthanized if their tumors exceeded 16 mm in diameter, became perforated, or if a mouse lost more than 20% body weight.

### Statistics and bioinformatics

Statistical analyses were conducted using GraphPad Prism version 10.3.1. Data are presented as individual data points and as mean ± SD or mean ± SEM. Statistical significance was evaluated using an unpaired two-tailed t-test, one-way, and two-way analysis of variance (ANOVA). A p-value of less than 0.05 was considered statistically significant. Survival differences in mice were assessed by the log-rank test. cBioPortal for Cancer Genomics was used to mine the TCGA data. Correlations between gene expression and NK cell infiltration were evaluated by Spearman’s rank correlation using TIMER2.0 (CIBERSORT-ABS algorithm).

## Acknowledgments

We thank Dr. Taru Muranen’s Lab from the Department of Medicine at Beth Israel Deaconess Medical Center, Harvard Medical School, for providing the patient-derived organoids. This work was supported by the Pelotonia Institute of Immuno-Oncology (PIIO). The content is solely the responsibility of the authors and does not necessarily represent the official views of the PIIO. We are also grateful to the Immune Monitoring and Discovery Platform (IMDP) of the PIIO at The Ohio State University Comprehensive Cancer Center (OSUCCC) for flow cytometry training. Special thanks to the former members of Vilgelm’s Lab, Marina Capece and Daniel de Lima Bellan, for their invaluable training and support.

## Author contributions

Conceptualization: AV, AM, DL, AF

Methodology: YW, NBK, ER, MSFP, DL, BO

Investigation: YW, ER, NBK, VB

Visualization: YW, ER, NBK, VB

Funding acquisition: AV, AM

Project administration: AV, YW

Supervision: AV, DL, AF

Writing – original draft: YW, AV

Writing – review & editing: all authors

